# The differentially expressed translationally controlled tumor proteins TCTP1 and TCTP2 in *Trypanosoma brucei*

**DOI:** 10.1101/190314

**Authors:** Borka Jojic, Simona Amodeo, Torsten Ochsenreiter

## Abstract

**Summary:** In *Trypanosoma brucei* we identified two TCTP genes differentially expressed during the parasite life cycle and discovered the mechanism by which this expression is controlled. Furthermore, we demonstrate that TCTP is important for cell growth as well as proper cell and organelle morphology in the insect form of the parasite.

**Abstract:** TCTP is a highly conserved protein ubiquitously expressed in eukaryotes. Studies have reported TCTP to be involved in growth and development, cell cycle progression, protection against cellular stresses and apoptosis, indicating the multifunctional role of the protein. Here, for the first time we characterize the expression and function of TCTP in the unicellular parasite *Trypanosoma brucei*. We identified two paralogue TCTP genes, which we named TbTCTP1 and TbTCTP2. They have identical 5’UTRs and only ten single nucleotide polymorphisms in the open reading frames (ORFs). However, the 3’UTRs differ dramatically in sequence and length. We found that the two TCTP mRNAs are differentially expressed during *T. brucei* life cycle. While procyclic form trypanosomes (PCF) express TCTP1, the bloodstream form trypanosomes (BSF), express TCTP2. We link the differential expression to the distinct 3’UTRs of the paralogues. In PCF cells, the protein appears to localize in the cytosol. We show that TCTP1 is essential for normal cell growth and has pleiotropic effects on the cells including aberrant cell morphology, enlarged and reduced number of acidocalcisomes and appearance of accumulations in the mitochondria.

## 3 Introduction

Since the time of its discovery about 30 years ago (Chitpatima et al., 1988), TCTP has continuously attracted research interest due to the high conservation among eukaryotes and involvement in a large number of biological processes (Bommer and Thiele, 2004). TCTP is expressed in most eukaryotes and in contradiction with the assigned name, not exclusively in cancerous tissues but in all the tested animal and plant tissues with preference towards the mitotically active ones and less in the post-mitotic tissues such as the brain (Berkowitz et al., 2008; Hinojosa-Moya et al., 2008; Sanchez et al., 1997; Thiele et al., 2000). In humans and rabbits the gene encoding for TCTP is transcribed in two mRNAs (named TCTP mRNA1 and TCTP mRNA2) which contain the same 5’UTR and open reading frame but different lengths of their 3’UTRs generated by alternative polyadenylation (Thiele et al., 2000). The two mRNAs are co-expressed but they differ in the ratio of their expression, with the shorter mRNA1 being more abundant. The functional significance of the existence of these two transcripts has not yet been identified. The predicted secondary structure elements of TCTP consist of three alpha-helices and eleven beta-stands and so far a microtubule binding, a Ca^++^ binding and two TCTP signature domains (TCTP1 and TCTP2) have been mapped (Bommer and Thiele, 2004). Despite the many years and number of research, an exact molecular function of TCTP has not yet been elucidated in any of the studied organisms. However, the different studies have related TCTP to many biological processes depending on the type of the cells/tissue, most notably growth and development, apoptosis, protection against cellular stresses and cell cycle (Berkowitz et al., 2008; Cao et al., 2010; Chan et al., 2012b; Chen et al., 2007; Gnanasekar et al., 2009; Gnanasekar and Ramaswamy, 2007; Hsu et al., 2007; Mak et al., 2001). Moreover several interacting/binding partners such as elongation factor eEF-1δ (Langdon et al., 2004), tubulin (Tuynder et al., 2002), calcium (Haghighat and Ruben, 1992) or Na, K-ATPase (Jung et al., 2004) have been identified. The protein is primarily localized in the cytosol, however in mammals and yeast it has been shown that TCTP can localize to the nucleus or mitochondria respectively, when cells are exposed to certain stress conditions (Diraison et al., 2011; Rid et al., 2010; Rinnerthaler et al., 2006). Here, for the first time we study the expression and function of TCTP in the unicellular parasite, *Trypanosoma brucei*. Trypanosomes are kinetoplastids, a group of flagellated protozoa with a major distinguishing feature being the presence of the kinetoplast (Stuart et al., 2008; Vickerman, 1976). They alternate their life cycle between the bloodstream of a mammalian host and different compartments of the tsetse fly *Glossina spp*. and undergo several differentiation steps in order to ensure its survival in the different environments (Vickerman, 1965). The differentiation steps are accompanied by extensive gene regulation that enable the parasite survival in hosts environments characterized among others by different energy sources, temperature and pH. Due to the continuous polycistronic transcription, unlike other eukaryotes, the regulation of individual gene expression occurs mainly at the post transcriptional level by *cis* and *trans-acting* elements (Clayton, 2013; Haile and Papadopoulou, 2007). The cis-acting elements are mainly motifs within the genes 3’UTRs, which confer repression, instability or increased translation of the mRNAs during parasites transition from one form to another (Hehl et al., 1994; Hotz et al., 1997; MacGregor and Matthews, 2012). As an adaptation to ensure host survival, not all the life cycle forms of *T. brucei* are proliferative, only the slender in the bloodstream of the mammalian host and the procyclics in the insect vector fly. The cell cycle is strongly connected to the parasites asymmetrical longitudinal cell morphology, maintained by a sub-pellicular microtubule corset (Hemphill et al., 1991). The microtubules in the corset are interconnected with each other and the plasma membrane and display a intracellular polarity with their plus ends at the posterior of the cell (Hemphill et al., 1991; Robinson et al., 1995). During cell division, the microtubular corset does not break down. Instead the newly synthesized microtubules are integrated into the old microtubule array making its inheritance between the daughter cells semiconservative (Sherwin and Gull, 1989). A number of single copy organelles and structures with a precisely defined cellular position are present in trypanosomes i.e. the flagellum, the nucleus, the kinetoplast, the mitochondrion, the basal body and the Golgi apparatus. During the cell cycle their duplication follows a specific temporal and spatial order to allow for proper cell division (McKean, 2003). Similarly to other eukaryotes, the cell cycle pattern in trypanosomes follows a G1, S, G2, mitosis (M) and cytokinesis phase (Hammarton, 2007; Woodward and Gull, 1990). However, different from animals, where cytokinesis requires the formation of an actomyosin contractile ring in trypanosomes the presence of actin is not required for cytokinesis and it rather depends on the proper separation and formation of the microtubule corset. (García-Salcedo et al., 2004). Moreover the placement of the cleavage site is asymmetrical (Wheeler et al., 2013). There are two important consequences of this asymmetry: (1) different zones of the dividing cell body undergo different growth and morphogenetic events and (2) the resulting two daughter cells inherit different portions of the existing cell body. One daughter cell receives the posterior part of the mother cell with the new flagellum, while the other daughter inherits the anterior part with the old-flagellum. The middle zone produces a novel anterior end for the new-flagellum daughter cell and a new posterior end for the old-flagellum daughter cell (Wheeler et al., 2013). Prior to this study the only existing information about TCTP in *T. brucei* or other kinetoplastids came (i) from a phylogeny study showing that despite being distantly related to the other commonly studies metazoans, TCTP in *T. brucei* is conserved (Hinojosa-Moya et al., 2008) and (ii) that it is a calcium binding protein (Haghighat and Ruben, 1992). Here we present a study characterizing the differential expression of two TCTP paralogues in *T. brucei*. We furthermore identify the mechanism by which the differential expression is controlled and characterize localization and loss of function phenotypes.

## 4 Results

### 4.1 Bioinformatics analysis of TbTCTP

We have identified a paralog of the previously described TCTP homolog in *T. brucei* (Hinojosa-Moya et al., 2008) The two genes are tandemly arrayed on chromosome eight and we named them TbTCTP1 (Tb927.8.6750) and TbTCTP2 (Tb927.8.6760). Phylogenetic analysis of the TCTP protein sequence confirms the very conserved primary structure throughout the eukaryotic supergroups (Fig S1) (Hinojosa-Moya et al., 2008). However, in contrast to previous reports most of the currently sequenced Kinetoplastea genomes contain two paralog TCTP genes similar to what has been described in Arabidopsis and Dictyostelium (Hinojosa-Moya et al., 2008). Within the Kinetoplastea the TCTP orthologues show up to 80% sequence similarity, while it exceeds 95% in the paralogs of this group. In several Kinetoplastea including *Leishmania major* and *Crithidia fasciculata* the paralogs are identical in the amino acid sequence. In *T. brucei* the paralogs contain five amino acid changes of which three are non-conservative (p. L23V, G54A, D62A, G64E, G85N; Fig 1A). The TCTP sequence conservation between Kinetoplastea and other Eukaryotes is up to 35% and includes the proposed microtubule, Ca^2+^ binding and the TCTP domains (Fig S2). Both genes have an identical 5’UTR and ten nucleotide changes in the ORF leading to the five changes at the amino acid level. However, the 3’UTRs of TbTCTP1 and TbTCTP2 differ drastically in sequence and length (Fig 1B).

**Fig 1.**
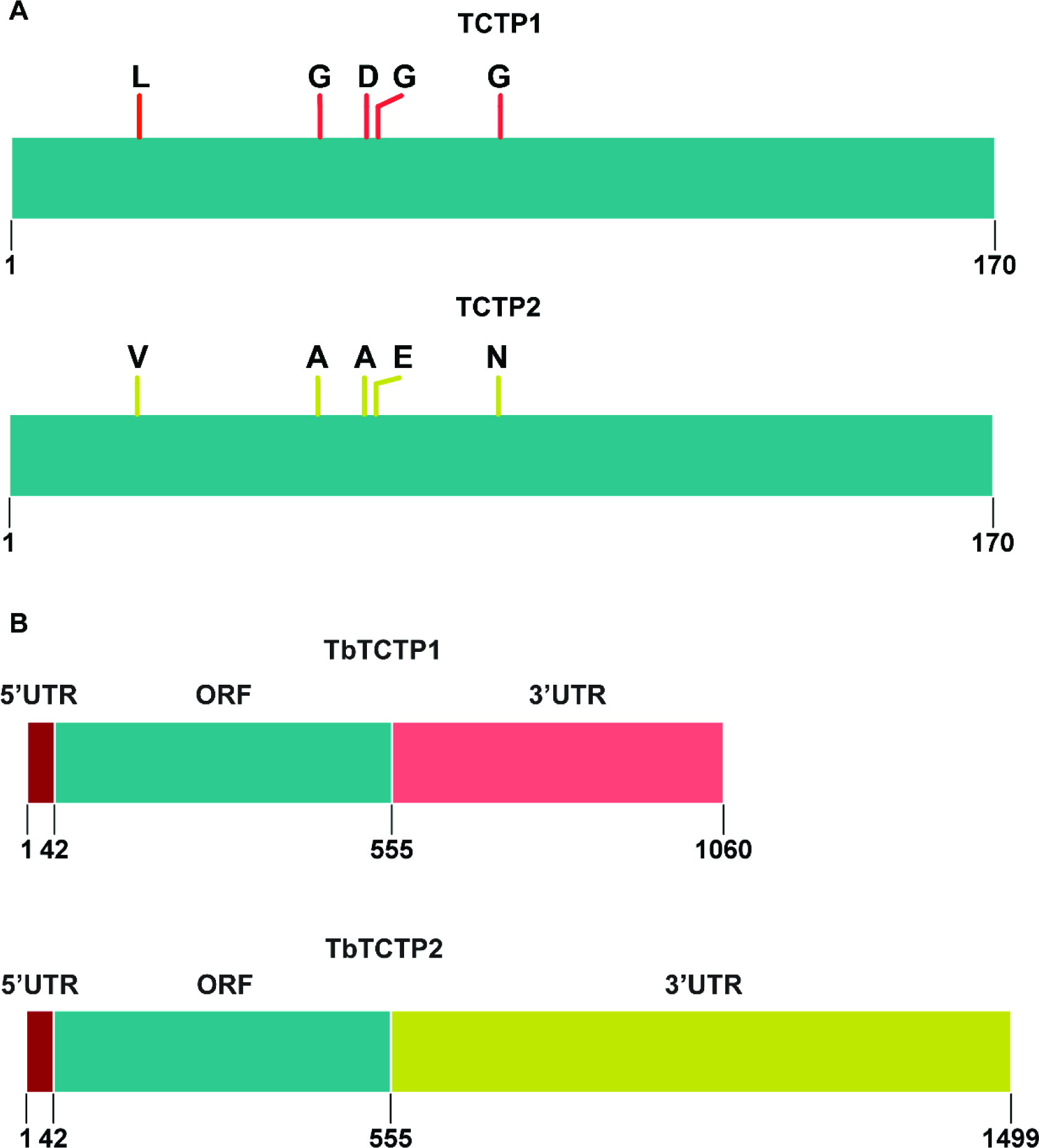
TCTP1 and TCTP2 in *T. brucei*. (A) Depiction of the TCTP1 and TCTP2 amino acid sequence. The proteins differ in only 5 amino acids at the positions 23, 54, 62, 64, 85 (B) Representation of the two TbTCTP paralogue genes in *T. brucei* (TbTCTP1 and TbTCTP2). Depicted are the 5UTRs (dark red), the open reading frame (ORF, light blue) and the 3'UTRs (light red, light green). The 5UTRs are identical in sequence.

### 4.2 TbTCTP paralogues expression in different life cycle stages of *T. brucei*

When we probed for TCTP expression in BSF and PCF parasites, we found that apparently two different isoforms of the gene were expressed (Fig 2A). We wondered if these isoforms represent the different paralogues and probed for the specific TCTP1 and TCTP2 3’UTRs in the BSF and PCF parasites. Northern blot analysis confirmed that TCTP1 mRNA is expressed in PCF trypanosomes while the TCTP2 mRNA is barely detectable in this life cycle stage (Fig 2B). The opposite is observed for BSF trypanosomes where TCTP2 is the predominantly expressed paralog (Fig 2C). Furthermore, we determined the half-live of each transcript in the life cycle stage where it is expressed. For this we incubated the cells with Actinomycin D (10 μg/ml) to stop transcription. Total RNA was collected at the indicated time points after Actinomycin D treatment (Fig 2B, C) and analyzed by northern blot. 18S ribosomal RNA was used as loading control. The mean relative mRNA abundance of TCTP1 and TCTP2 from eight time points following Actinomycin D treatment was measured and showed a continuous decrease. The half-live of TCTP1 mRNA in PCF was calculated to be ±217 minutes while the half-live of TCTP2 mRNA in BSF was ±51 minutes (Fig 2D). To better understand the mechanism of stage specific expression and the role of the 3’UTRs we created two constructs in which the TCTP1 or TCTP2 3’UTR is linked to a reporter gene (chloramphenicol acetyltransferase, CAT, Fig 3A). Both constructs were transfected in PCF and BSF trypanosomes to integrate into the tubulin locus. Total RNA from three different clones of each life cycle stage was isolated and further analyzed by probing against the CAT ORF (Fig 3B, C). We observed that in PCF trypanosomes the CAT-TCTP1 construct was well expressed in all clones, while the CAT-TCTP2 construct was barely detectable (Fig 3C). In the BSF cells, the CAT-TCTP2 was detected in all clones whereas the CAT-TCTP1 construct was not detectable (Fig 3B). Thus, the 3’UTRs of TCTP1/2 are responsible for the differential expression of TCTP1 and TCTP2 in PCF and BSF trypanosomes, respectively.

**Fig 2.**
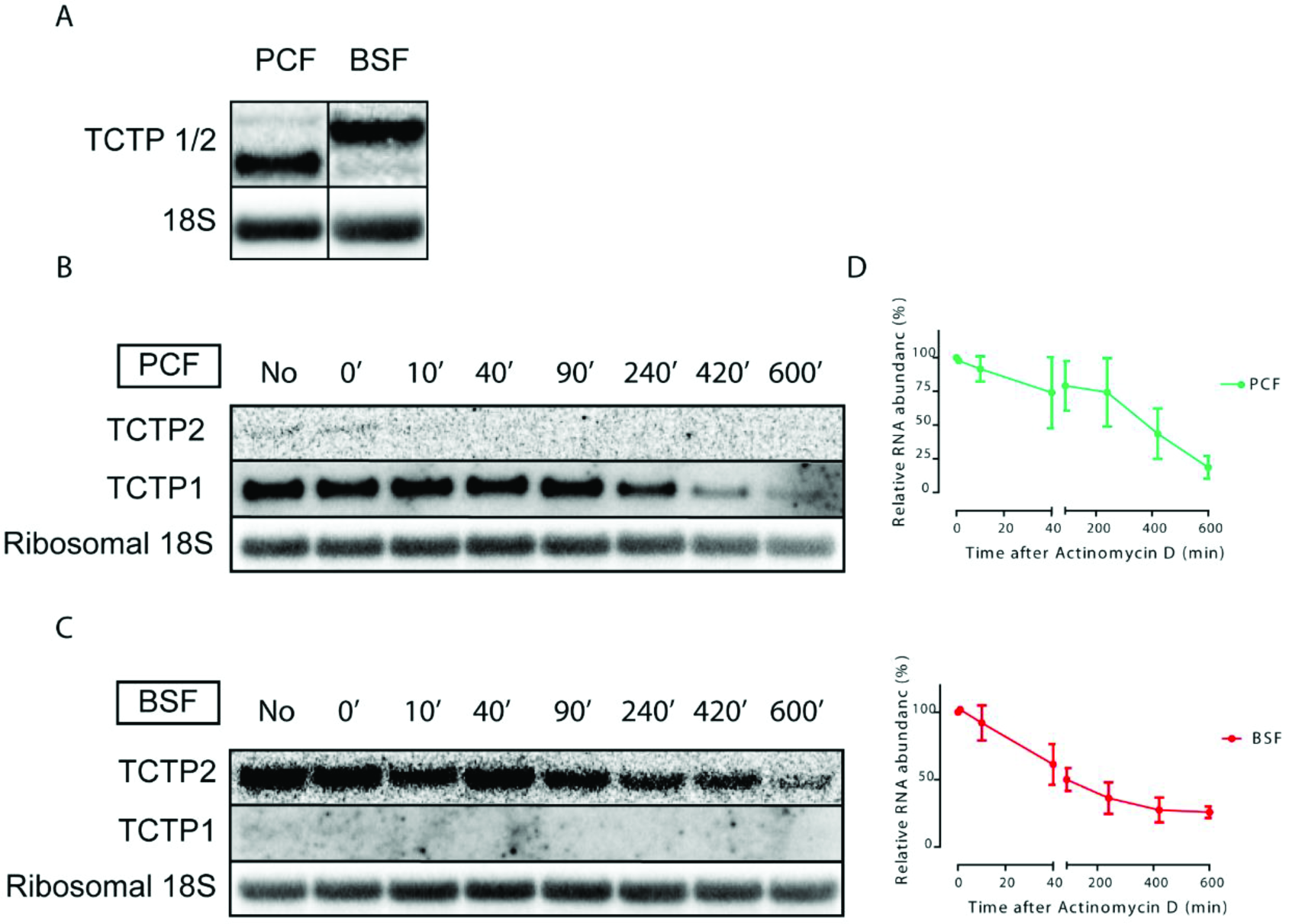
TCTP mRNA expression is developmentally regulated in *T. brucei*. (A) TCTP mRNA expression in BSF and PCF trypanosomes. (B-C) Northern blot analysis of TCTP1 and TCTP2 mRNA from BSF and PCF trypanosomes. The RNA was collected from cultures incubated with Actinomycine D (10 μg/ml) at the indicated time points (minutes). Ribosomal 18S RNA is used as a loading control. (D) Quantification of the relative RNA abundance of TCTP1 in procyclic and TCTP2 in bloodstream trypanosomes. Three independent experiments were performed for each timepoint. Error bars represent standard deviation. Values are normalized to 18S rRNA. mRNA abundance in untreated cells was assigned 100%. Half-lives were calculated to be 217 min in PCF and 51 min in BSF.

**Fig 3.**
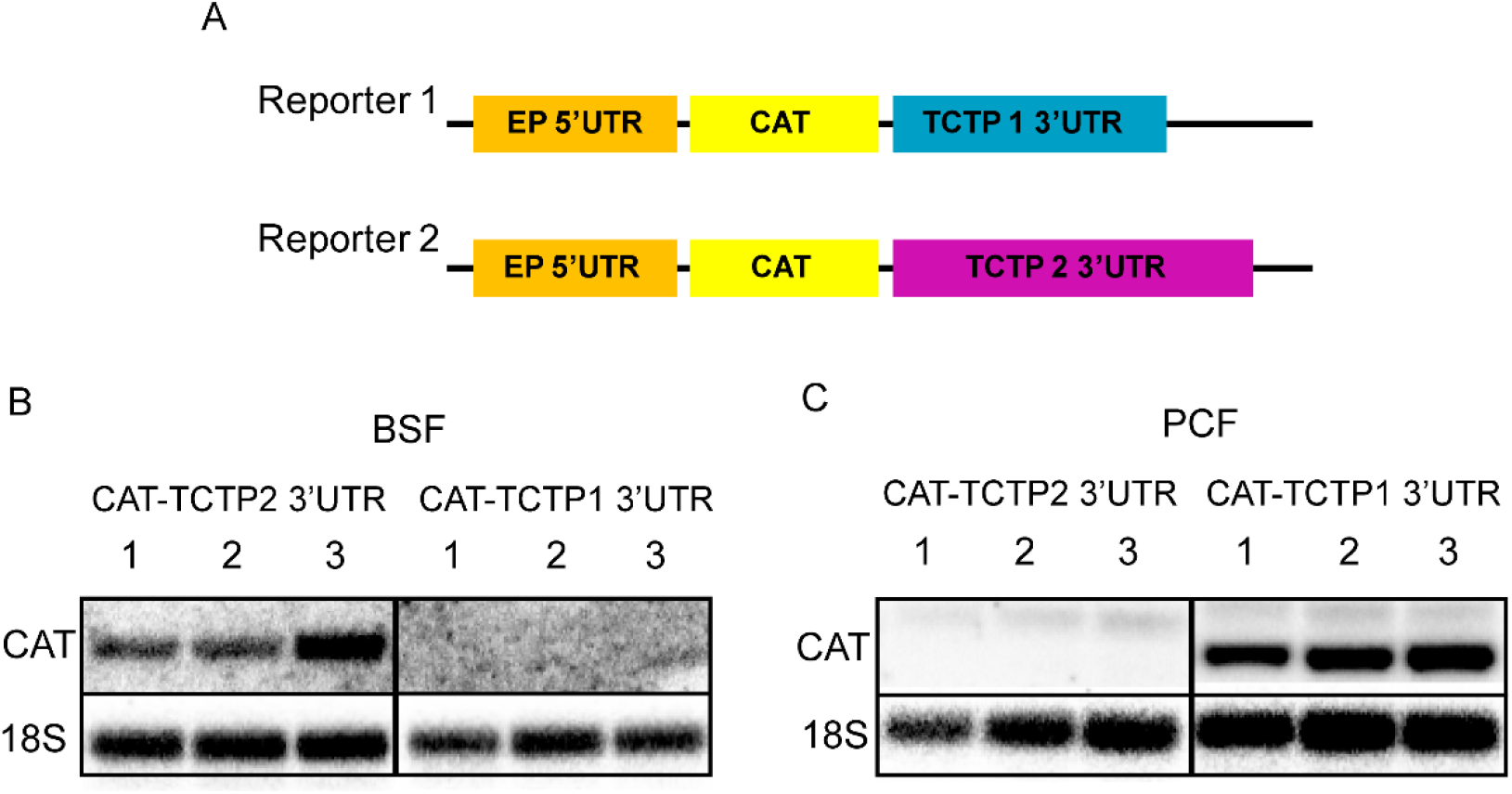
Role of the distinct TCTP 3’UTRs in life stage specific mRNA expression. (A) Schematic representation of the two reporter constructs linking the EP 5’UTR (orange) and CAT ORF (yellow) to the TCTP1 (blue) or TCTP2 (purple) 3’UTR. Each construct was transfected in BSF and PCF trypanosomes. (B-C) Total RNA from three clones per condition was isolated and CAT mRNA expression was analysed by northern blot probing for CAT ORF. Ribosomal 18S was used as a loading control.

### 4.3 Localization of TCTP1 in PCF trypanosomes

For the further analysis of the localization and function, we focused on TCTP1 in the PCF cells. In order to localize TCTP1 we produced a polyclonal antibody in rats. In biochemical fractionations of total cell extracts with digitonin TCTP1 localized to the cytoplasmic fraction (Fig 4). Since the antibody did not work in immunofluorescence microscopy, we tagged TCTP1 N-(myc) and C-terminally (triple HA) and checked for its localization using anti-myc and anti-HA antibodies. The ectopically expressed TCTP1 was distributed in the cell in a pattern that is consistent with a cytoplasmic localization. Additionally, we found a discernible depletion of TCTP in the nucleus (Fig 5), and did not detect any change of localization or abundance during the cell cycle (Fig 5). The *in-situ* c-terminally tagged TCTP1 showed the same localization pattern (Fig S3).

**Fig 4.**
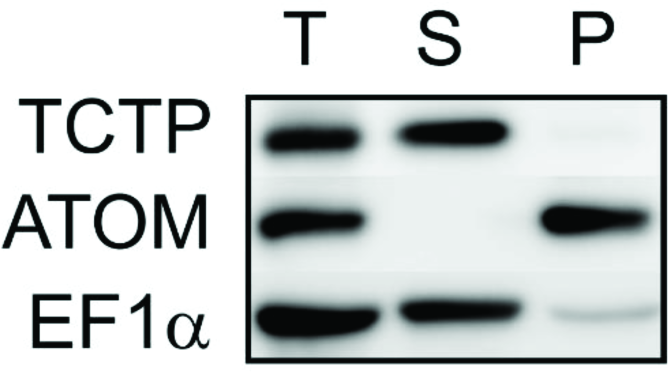
TCTP localizes in the cytosolic fraction of the cells. Procyclic form trypanosomes were lysed with 0.025% digitonin and the cell extracts were fractionated by differential centrifugation. Total cellular extract (T), supernatant (S) and pellet (P) fractions were analysed by western blot decorated with antibodies against TCTP, ATOM and EF1a.

**Fig 5.**
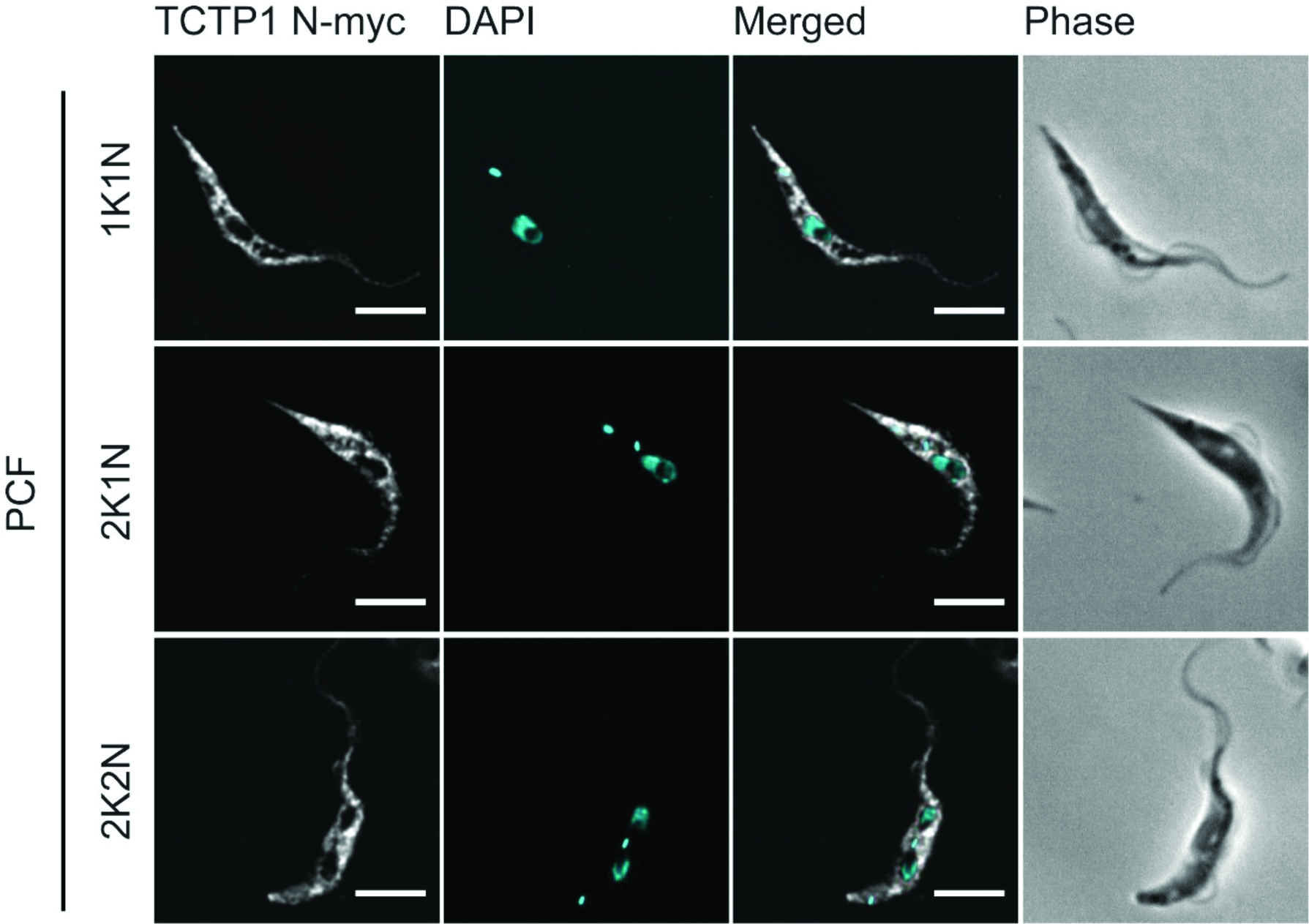
Localization of N-terminally tagged TCTP in procyclic trypanosomes. Inducible ectopic copies of N-terminally myc-tagged TCTP were expressed for 10 hours in procyclic trypanosomes. Proteins were detected by immunofluorescence microscopy with anti-myc antibody (white). The DNA is detected by DAPI (cyan). Scale bars: 5 μm

### 4.4 Depletion of TCTP1 leads to a slow growth phenotype and affects several organelles and cytokinesis

To determine the function of TCTP1 in PCF parasites we depleted TCTP mRNA by performing RNAi against the TCTP ORF. We could show that TCTP protein levels were strongly depleted after 48 hours of RNAi induction (Fig 6 inset). PCF cells started growing slower at day three post induction and continued to do so at least until day eight. We analyzed the karyotype and the morphology of the TCTP depleted cells by DAPI staining for the nucleus and kinetoplast DNA, and phase contrast images (Fig 7A). Upon eight days of TCTP RNAi induction we observed an increase on 1K1N cells from 71% to 81% and a decrease of 2K1N cell from 21% to 12% (Fig 7B). We noticed an accumulation of cells displaying a “tadpole” morphology (up to 40%, Fig 7C). These cells are characterized by a distinct shape of the cell body where the posterior end of the cell is round and enlarged whereas the anterior becomes more slender than in the wild type situation (Fig 7A). This change in morphology was confirmed by calculating and comparing the ratio of the cell body length (L) to the cell body diameter through the nucleus (D) as well the cell surface area in the same population of non-induced (n = 20) and induced cells (n = 24). While there was no significant change in the mean cell surface area of the induced population (Fig 7E), a significant reduction in L/D ratio was calculated for the cells where TCTP was depleted (Fig 7D). When we measured tadpoles only (n = 7), the mean surface area is reduced from an average of 40 μm^2^ to an average of 29 μm^2^. The L/D ratio was also reduced from an average of eight in the non-induced population to an average of 5.4 in the induced ones. This phenotype was observed exclusively in 1K1N cells indicating that the change in shape is likely a consequence of improper cytokinesis.

**Fig 6.**
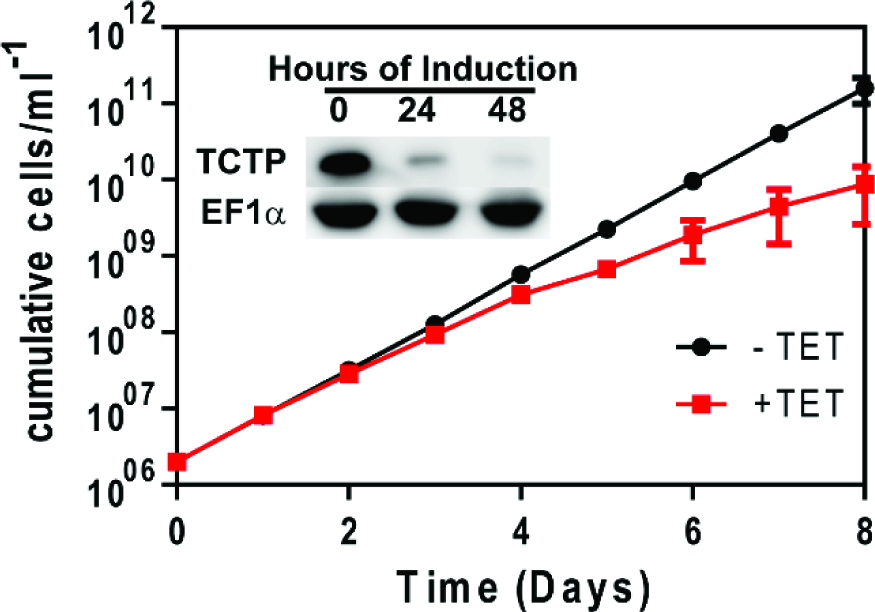
The effect of TCTP downregulation on the cell growth. Growth of cells with or without addition of tetracycline was monitored for 8 days. Inset: western blot showing TCTP downregulation upon 24 and 48 hours of RNAi induction. EF1a is used as loading control.

**Fig 7.**
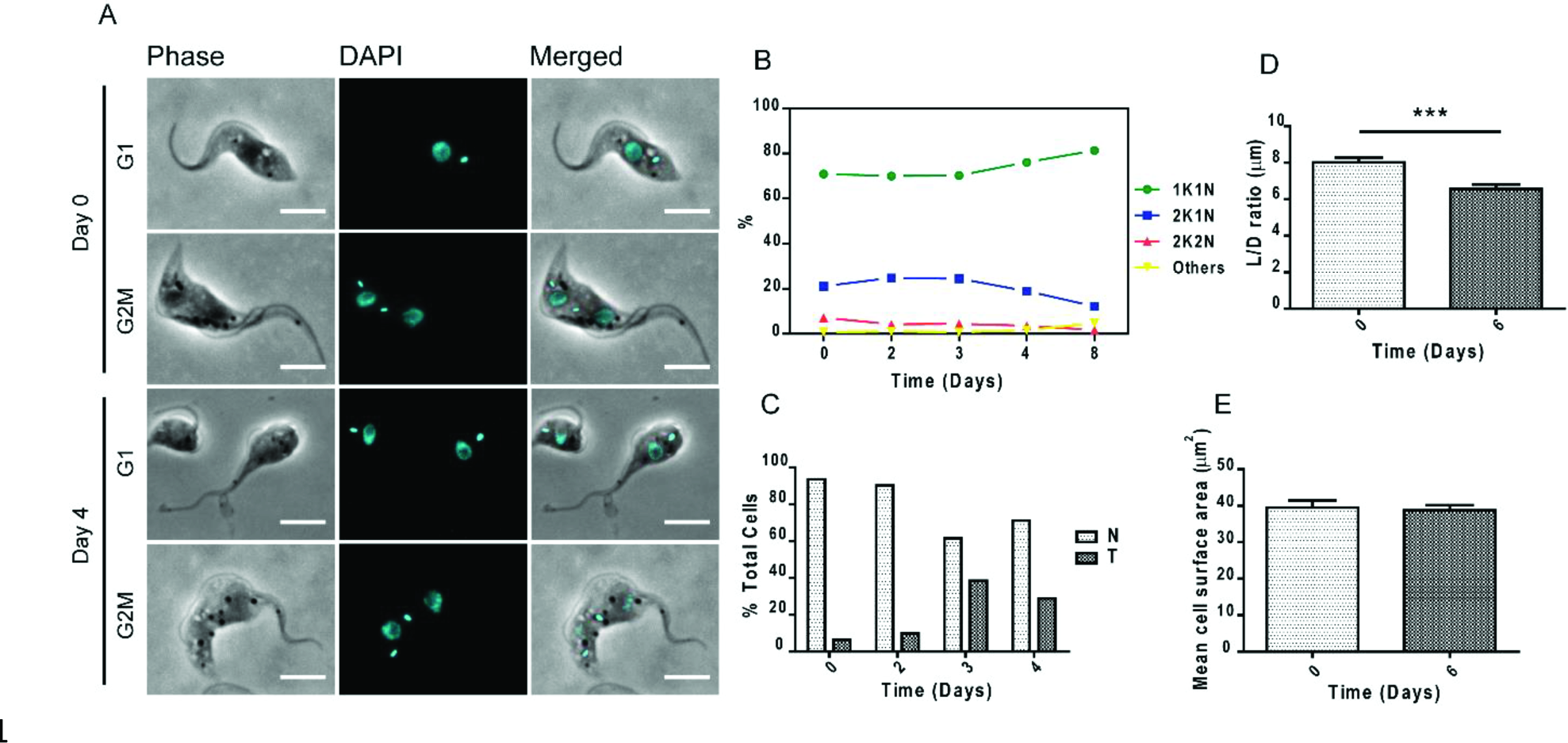
The effect of TCTP downregulation on the morphology of procyclic trypanosomes. (A) Representative single cell images of non-induced and tadpole-like trypanosomes in G1 and G2M stages of the cell cycle from a population of day 4 induced TCTP RNAi trypanosomes (B) Percentage of cells in different cell cycle stages and (C) percentage of cells having the tadpole phenotype in a population of TCTP RNAi procyclic trypanosomes (n ≥ 250 for each timepoint). (D) Cell body length (L) to cell body diameter (D) ratio of cells from a population of non-induced and induced (day 6) TCTP RNAi (***p=0.0004, Two-tailed student’s t-test, n ≥ 20). (E) Same cell populations as in (D) were used to measure cell surface area change upon TCTP RNAi. Scale bars: 5 μm.

Since TCTP has been shown to be a Ca^2+^ binding protein in trypanosomes (Haghighat and Ruben, 1992) we wondered if depletion of the protein has an effect on the acidocalcisomes (ACs), the major Ca^2+^ storage organelles in the cells. We visualized the ACs by immunofluorescence microscopy using the vacuolar proton pyrophosphatase (VP1) antibody (Fig 8A) (Rodrigues et al., 1999). Additionally, the DNA and cell morphology were visualized by DAPI and phase contrast images (Fig 8A). We noticed that upon six days of TCTP downregulation the ACs were enlarged in size and the VP1 visualized no longer as small dots but as larger ring-like structures (Fig 8A). This change in ACs morphology occurred in all cell cycle stages. We visualized and confirmed this difference by images acquired with super resolution microscopy (Fig 8B). We counted the total number of ACs in non-induced and induced cells (n=10) and within each cell sorted them as dot-like or ring-like shaped. We noticed that in induced cells not only the total number of ACs per cell is reduced from an average of 21 to an average of 14, but also the number of ring-like ACs per cell increases up to four-fold (Fig 8D). Furthermore, we looked at the ultrastructure of the acidocalcisomes before and after TCTP depletion by transmission electron microscopy (TEM). As previously shown ACs are recognized as vesicles either empty or containing electron dense material (Fig 8C) (Rodrigues et al., 1999). The mean area of the acidocalcisomes was measured in random thin slices (n=80) from RNAi induced and non-induced cells. As already seen in the fluorescence microscopy the size of the acidocalcisomes increases significantly (Fig 8E). We also tested if the organelle size increase might correlate with an increased storage of phosphate in the organelles however were unable to identify any differences by epifluorescence microscopy. The second most important storage compartment for calcium is the mitochondrion. In order to check if the depletion of TCTP leads to mitochondrial structure abnormalities we visualized the PCF mitochondria by immunofluorescence microscopy using an antibody targeting the mitochondrial matrix heat shock protein 70 (HSP-70). The images showed that upon six days of TCTP depletion, accumulations appeared within the mitochondrial network (Fig 9A, arrowheads point to accumulations). In the vast majority of cases, there was a single accumulation within one cell and it is located predominantly between the nucleus and kDNA. As shown in Fig 9A, formation of these mitochondrial accumulations can occur in all stages of the cell cycle. The number of accumulations increased from 10% in the non-induced cells (n=160) to 40% in the TCTP RNAi induced cells (n=160) (Fig 9B).

**Fig 8.**
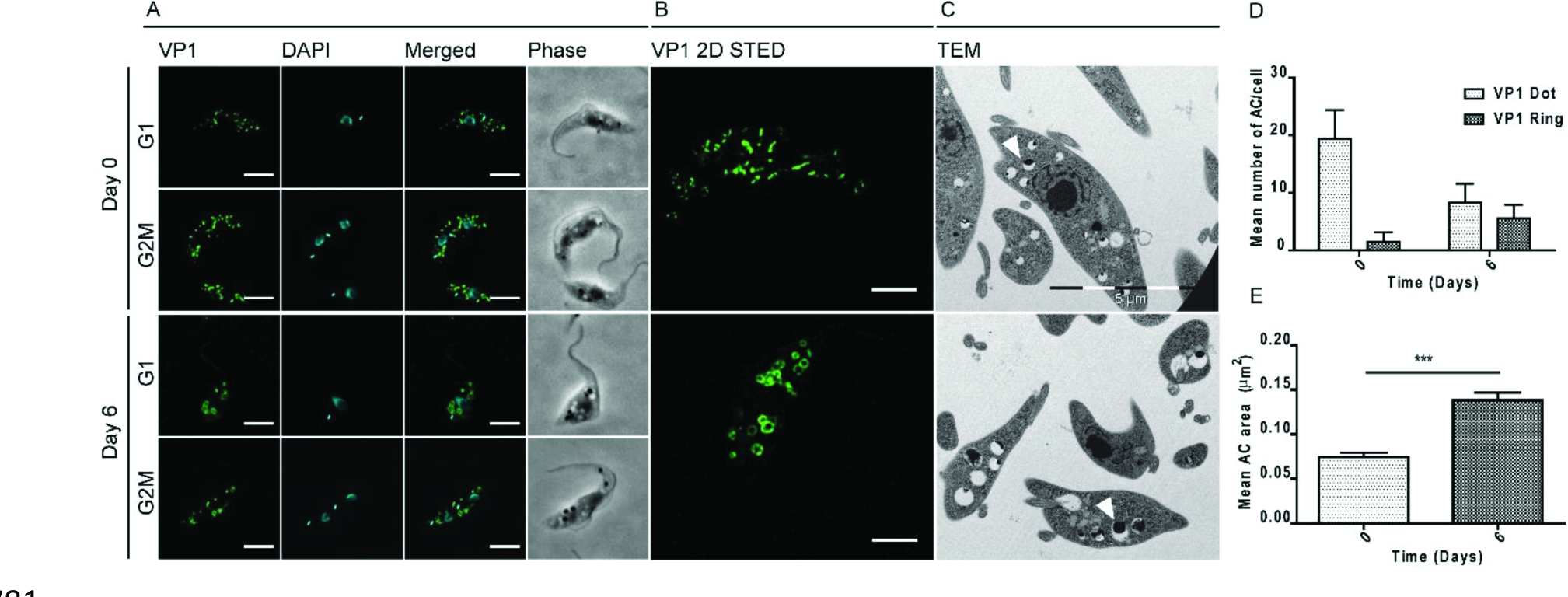
Morphology of acidocalcisomes following downregulation of TCTP. Non-induced and induced (Day 6) PCF trypanosomes were stained for (A) acidocalcisome marker VP1 (green) and DAPI (cyan) for the nucleus and kDNA. Cell morphology is shown by phase contrast images (gray). (B) Representative images of acidocalcisomes in single cells acquired by super resolution microscopy. (C) Representative images of acidocalcisomes ultrastructure by transmission electron microscopy. Arrowheads point to examples of acidocalcisomes size in non-induced and induced cells. (D) Histograms showing the mean number of dot-like versus ring-like acidocalcisomes per cell in D0 and D6 cells (n=10) (E) Quantification of mean acidocalcisome area in TEM sections of D0 and D6 cells (n=80, one tailed t-test, p<0.0001). Bars represent SEM). Scale bars: (A) 5 μm, (B) 2 μm.

**Fig 9.**
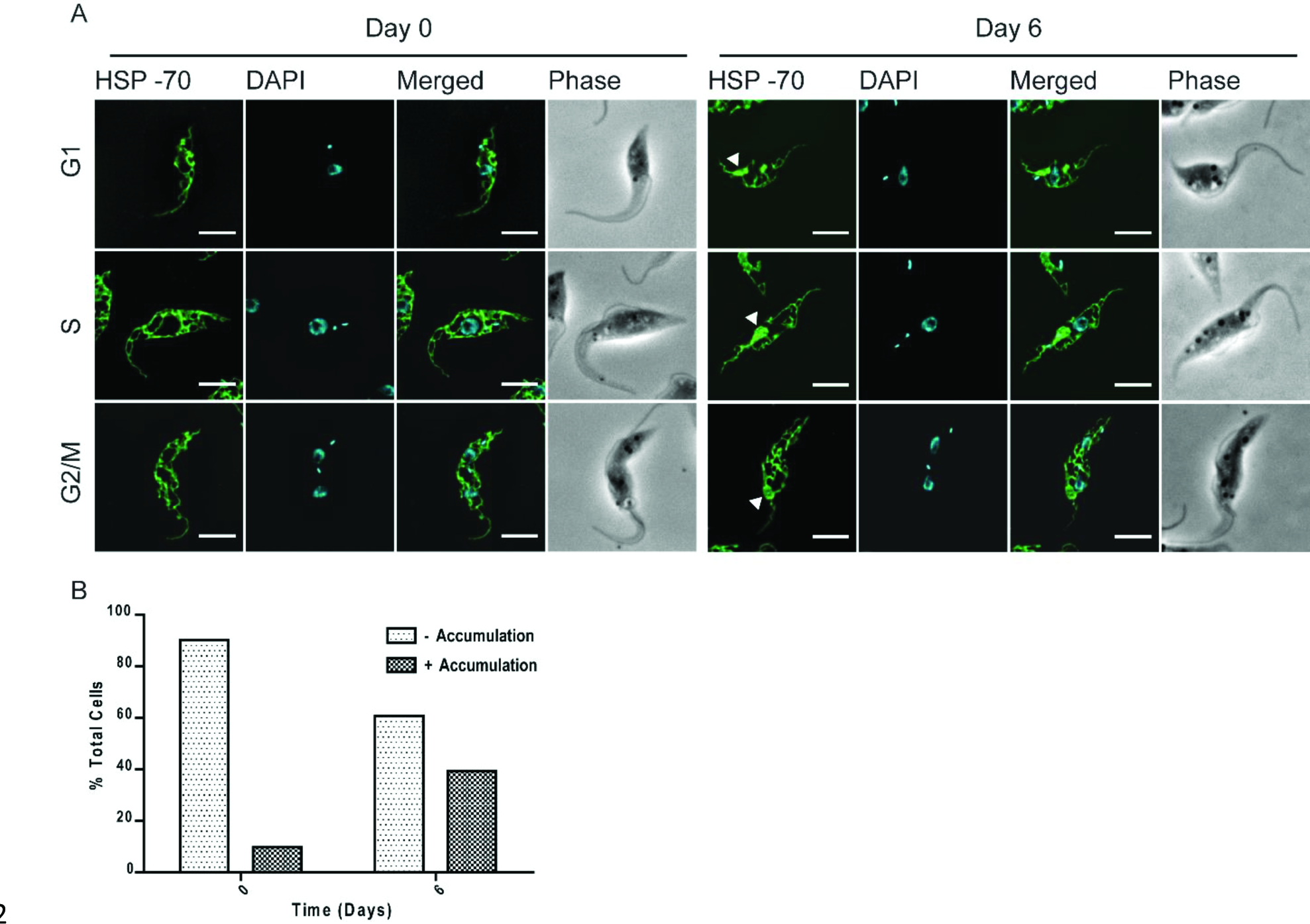
Effect of TCTP downregulation on mitochondria. (A) Immunofluorescence images of non-induced (Day 0) and induced (Day 6) procyclic cells stained for mitochondria (HSP-70, green) and DAPI (nucleus, kDNA, cyan) are shown. Phase contrast images (grey) show cell morphology. Arrowheads point to mitochondrial accumulations. Representative images of cells in G1, S and G2M cell cycle stages are selected for both non-induced and induced cell populations. (B) Quantification of mitochondrial accumulations in D0 and D6 induced cells (n=160). Scale bar: 5 μm.

## 5. Discussion

TCTP has gained significant attention in recent years since it has been recognized as therapeutic target in prostate, breast and lung cancers. Consequently, more than 300 publications describe the large variety of processes TCTP might be involved in, including apoptosis, cell cycle regulation, stress response, just to name a few (Chan et al., 2012a). Surprisingly little is known about the molecular mechanism by which TCTP is involved in these many different processes or how the gene might be regulated.

This is the first report about localization, function and regulation of TCTP in the eukaryotic super-group of the Excavates. In addition to the annotated TCTP gene, in *T. brucei* we identified a paralog to be encoded in tandem. This feature seems to be conserved within the Trypanosomatida while *Bodo saltans* and *Perkinsela sp*. for example only to contain one orthologue in the current genome annotation (BS72265, *Bodo saltans*. KNH04185, *Perkinsela* sp.). This, together with the fact that the sequence conservation of the two TCTP paralogues within the different Trypanosomatida is very high might point towards a duplication event just prior to the separation of the Trypanosomatida from the other Kinetoplastea.

While the 5′ UTRs of two paralogues in *T. brucei* are identical and the ORFs only contain ten nucleotide polymorphisms, the 3′ UTRs are very different in length and sequence. A similar situation has recently been described for two genes encoding cytoskeletal proteins in *T. brucei*, however there the mechanism of gene expression regulation remained unexplored (Portman and Gull, 2014). For the two TCTP paralogues in *T. brucei* we provide strong evidence that the mechanism by which these two genes are differentially regulated is based on the different 3' UTR sequences providing a prominent example of posttranscriptional regulation in *T. brucei*. The difference in half-life of the two TCTP transcripts (Fig 2D), can at least partially be attributed to the different growth rates of the two life cycle stages. Interestingly, the feature of two different 3' UTRs in TCTP transcripts is also found in other species. In humans and rabbits for example the TCTP gene is transcribed as two mRNAs that only differ in the length of their 3’UTR as a result of differential polyadenylation (Thiele et al., 2000). Both mRNAs are co-expressed in almost all tissues. Interestingly, the TCTP transcript with the shorter UTR is often times more abundant than the long UTR counterpart, however so far no function has been assigned to this observation (Thiele et al., 2000). Thus, the different UTRs might also have a function independent of the protein product a possibility that has previously been described albeit for the entire TCTP mRNA. Bommer and colleagues suggest that the TCTP mRNA is a highly folded mRNA that can bind to and activate the dsRNA-dependent protein kinase (PKR) an important regulator of translation (Bommer et al., 2002). Thus, one could speculate that the different TCTP 3' UTRs might also be involved in the specific biology of the two different life cycle stages.

The cytoplasmic localization of TCTP1 in *T. brucei* is similar to what has been described for most organisms (Acunzo et al., 2014; Rinnerthaler et al., 2006), and has recently been confirmed by a tagging screen of the TrypTag consortium on the TriTrypDB website. However, the TrypTag high throughput study additionally identified a localization at the flagellum, which we did not detect in epifluorescence microscopy or biochemical fractionations. We also did not detect any changes in the localization of TCTP during the cell cycle. As mentioned above a large number of functions are ascribed to TCTP in the different model systems, and very little is known about the mechanisms by which the functions are performed. We used RNAi and efficiently depleted the TCTP mRNA in procyclic trypanosomes. While TCTP depletion led to a growth defect, it seemed to not affect all cells in the population. After several days of RNAi induction up to 40% of cells showed a distinct tadpole like morphology phenotype the majority of the cells had wild type appearance. It is tempting to speculate that the tadpole shaped cells are a dead end and cannot further divide and thus are the reason for the decrease in growth rate of the population. Future experiments focusing on cells in division will address this question.

During cytokinesis of PCF parasites the anterior part of the cell including the new flagellum is inherited to one-while the posterior region with the old flagellum is inherited to the other daughter cell. Consequently the central part of the cell has to be remodeled such it can give rise to a new posterior for the daughter cell with the old flagellum and a new anterior for daughter with the new flagellum (Wheeler et al., 2013). We hypothesize that this remodeling process and more specifically the posterior end processing of the new daughter cell with the old flagellum is impaired through the loss of TCTP1. The involvement of TCTP1 in this process is supported by the findings in other systems where TCTP has been demonstrated to be a microtubule binding protein and to be involved in the formation of the mitotic spindle (Gachet et al., 1999). Interestingly we also detected a change in mitotic spindle staining in the PCF cells upon RNAi (Fig S4) that would support the potential role of TCTP1 as a tubulin binding protein. While at this point, we have not further experimental evidence for the tubulin interaction activity of TCTP1 we can already exclude that it is involved in all tubulin turnover/processing events since we did not detect any effect on the basal body or flagellum biogenesis in PCF TCTP RNAi knockdown cells (Fig S5-S6).

TCTP down regulation in PCF also had an effect on the morphology and the number of acidocalcisomes, the main calcium and polyphosphate storage organelles, which were first discovered in trypanosomes and later in other organisms (Docampo and Moreno, 2011; Moreno et al., 1992). Upon downregulation of TCTP we observed that the number of acidocalcisomes per cell decreased, while the size of the remaining organelles almost doubled. We tested if this size increase could be due to an accumulation of polyphosphates but did not detect any changes by epifluorescence microscopy. Future experiments should address the questions if and how TCTP1 is involved in Ca^2^+ homeostasis and how that is linked to the changes in mitochondrial and acidocalcisome morphology.

In conclusion, this study for the first time characterizes the universally conserved TCTP protein in the Excavates and attributes a function to the different 3’UTRs of TCTP, a feature that is seen in many different groups. Our experiments clearly demonstrate that the 3’UTRs are the responsible elements for the differential expression of TCTP1 and TCTP2 paralogues in the BSF and PCF trypanosomes. Similar to what has been described previously in other groups TCTP seems to be involved in a variety of functions and based on the acidocalcisome and mitochondrial phenotypes we add Ca^2^+ homeostasis to the list of potential TCTP entanglements.

## 6 Materials and methods

### 6.1 Trypanosome cell lines and culturing

For RNAi and gene tagging experiments we used transgenic *T. brucei* bloodstream (New York single marker, NYsm) and procyclic (29-13) cell lines co-expressing T7 RNA polymerase and a tetracycline repressor (E Wirtz et al., 1999). The BSF cells were cultured at 37°C and 5% CO2 in HMI-9 medium supplemented with 10% FCS (Hirumi and Hirumi, 1989) in the presence of 2.5 μg/ml geneticin (G418). The PCF cells were maintained at 27°C in SDM-79 medium with 10% FCS (Brun and Schönenberger, 1979) in the presence of 15 μg/ml geneticin and 25 μg/ml hygromycin. For the CAT assay we used wild-type *T. brucei* 427 BSF expressing VSG221 (MITat 1.2) and wild-type 427 PCF. The cells lines were maintained in the same media conditions as the transgenic ones but in absence of the antibiotics. The cell lines were obtained from the established collection of the Institute of Cell Biology, University of Bern, Bern, Switzerland

### 6.2 Plasmid constructs and transfection

For inducible RNAi against TCTP1 and TCTP2 mRNAs we used a pLEW100 based stem-loop plasmid (Mani et al., 2015; Elizabeth Wirtz et al., 1999) where an insert of 512 bp targeting the full ORF sequence of the TCTP1 gene was integrated. The constructs were linearized with NotI and 10 μg were transfected in NYsm BSF and/or 29-13 PCF by electroporation. The positive clones were selected with blasticidin (5 μg/ml in BSF and 10 μg/ml in PCF). Induction of RNAi was done by addition of 1 μg/ml tetracycline. For the C-terminal tagging one allele of TCTP1 in PCF was *in situ* tagged with a triple Hemagglutinin (HA) epitope (Oberholzer et al., 2006). For inducible N-terminal c-Myc tagging, the full ORF plus the first 21 nt from the 3’UTR of TCTP1 were amplified by PCR and cloned in pJM-2 vector (a gift from A. Schneider; (Mani et al., 2015). Upon transfection (as described above) the clones were selected with puromycine. Expression was induced by addition of 1 μg/ml tetracycline. For the chloramphenicol acetyltransferase (CAT) activity assay the 3’UTR of TCTP1 (1-505 nt) or TCTP2 (1-944nt) was cloned into pHD2169 vector between BamHI and SalI restriction sites (a gift from C. Clayton). The plasmid was linearized by NotI digestion at the β-tubulin targeting sequence. The constructs were transfected in wild-type BSF and PCF cells (described above) and the clones were selected with G418.

### 6.3 Northern blotting and RNA analysis

For northern blot analysis, total RNA was extracted from trypanosomes by re-suspending 10^8^ trypanosomes in 1 ml RiboZol™ (Amresco). The RNA was purified by phenol/chloroform, precipitated with ethanol and unless directly used, stored at ‐20^0^C. 8-10 μg RNA was resolved in 1.4 *%* agarose gels and transferred overnight (ON) onto nylon cellulose membranes (Millipore) by capillary blotting. The membranes were incubated ON with radioactively labelled (α-P32-dCTP) PCR generated probes using Random Primed DNA Labeling Kit (Roche). For normalization the blots were re-probed for 18S ribosomal RNA (rRNA). The 18S rRNA probe was generated by T4 PNK labelling of an 18S oligonucleotide with γ-P32-ATP. The probes for the ORF and 3’UTRs of TCTP1 and TCTP2 were generated by amplifying the ORF or specific fragments of the 3’UTRs post the Stop codon: 14-488 nt for TCTP1 and 6-606 nt for TCTP2 using the genomic DNA as template. The CAT probes were generated by amplification of the CAT ORF (13-436 nt) using pHD2169 as template. For the mRNA decay analysis, the transcription was blocked by addition of Actinomycine D (10 μg/ml) and total RNA was isolated from the cells (as described above) at different time points after addition (0 min, 10 min, 40 min, 90 min, 240 min, 420 min and 600 min). Three biological replicates were performed. GraphPad Prism was used to plot the mean relative mRNA abundances at each time point (normalized to 18S RNA) and calculate the mRNA half-lives.

### 6.4 Generation of polyclonal TCTP antibody

The full length ORF of TCTP1/2 was cloned into pHIS-parallel-1 vector which allows the expression of TCTP1/2 as a fusion protein with the HIS_6_ peptide (Sheffield et al., 1999). Expression of the fusion protein was induced for 2 hours in *E. coli* BL21 paplac strain at OD_600_=0.8 by addition of 1 mM isopropyl β-D-1-thiogalactopyranoside (IPTG). Next, the bacteria were kept in cold, re-suspended in lysis buffer (50 mM NaH_2_PO_4_, 300mM NaCl, 5mM Imidazol, 10% v/v Glycerol, protease inhibitors, pH=8.00), sonicated and spun down at 9500g for 30 minutes at 4^0^C. The supernatant containing the proteins was loaded into Ni-charged IMAC resin columns (*BIO-RAD*) and the HIS_6_-TCTP1/2 fusion protein was purified by immobilized metal affinity chromatography in elution buffer (50 mM NaH_2_PO_4_, 300mM NaCl, 20mM Imidazol, pH=8.00) and used to produce rat polyclonal antibody (Eurogentec, Belgium).

### 6.5 Western blot

For western blotting trypanosome pellets were washed in phosphate buffered saline (PBS, pH=7.2) and re-suspended in standard Laemmle buffer (LB) (10^6^ in 15 μl). For the digitonin fractionation the cells were washed once in PBS, then the pellets were re-suspended in SoTE buffer (0.6 M sorbitol, 2 mM EDTA, 20 mM Tris-HCl, pH 7.5) containing 0.025% digitonin and incubated on ice for 5 min. Next, the cell fractions were separated by differential centrifugation at 8000 rcf for 5 min at 4°C. This gives a supernatant fraction enriched in cytosolic proteins and a pellet enriched in organelles and organelle bound proteins. Both fractions were lysed in LB, boiled for 5 min at 95^0^C, cooled on ice for 5 minutes and loaded on 10% or 12% SDS-polyacrylamide gels (10^6^-10^7^ cells per lane) before subjected to western blotting. Next the proteins were transferred onto PVDF Immobilon-P membranes (Millipore) using BioRad wet blotting system and blocked for 1 hour at room temperature in 10% skimmed milk or BSA solution in PBST (PBS + 0.1% TWEEN-20). The primary antibodies were incubated for one hour at room temperature or overnight (ON) at 4^0^C in PBST. In this study we used rat-polyclonal α-TCTP (1:50, Eurogentech); rabbit α-ATOM (1:10000, (Pusnik et al., 2011)); mouse α-EF1α (1:10000, SantaCruz). After washing the primary antibody by incubating the membranes 3 times, 10 minutes each time in PBST, the membranes were incubated for 1 hour at room temperature with the secondary antibody. Secondary antibodies were: rabbit α-rat HRP-conjugate (1:10000, Dako), rabbit α-mouse HRP-conjugate (1:10000, Dako) and swine α-rabbit HRP-conjugate (1:10000, Dako). For the chemiluminescent detection, the SuperSignal system (Pierce) was used and images were acquired with Amersham Imager 600.

### 6.6 Immunofluorescence and microscopy

For immunofluorescence, BSF or PCF cells were harvested by slow centrifugation (2000 rpm/5 min), washed once in PBS and then fixed on slides for 4 min with 4% PFA in PBS. The cells were permeabilized for 5 min with 0.2% TritonX-100 in PBS and blocked for 30 min with 4% BSA in PBS. The cells were incubated for 1 hour in a wet chamber with primary antibody diluted in 4% BSA in PBS, washed three times in PBST and incubated again in dark wet chambers with secondary antibodies diluted in 4% BSA in PBS. The cells were mounted with ProLong^®^ Gold Antifade Mountant with or without DAPI (Invitrogen). Images were acquired with Leica DM 5500 fluorescent light microscope and/or Leica SP8 Confocal Microscope System with STED and deconvoluted by Leica LAS AF and Huygens software, respectively. The primary antibodies used in this study were: rat α-PFR (1:1000, (Kohl et al., 1999)), rat α-YL1/2 (1:2000), mouse α-KMX (1:500, (Sasse and Gull, 1988)), mouse α-mtHSP70 (1:2000, (Klein et al., 1995)), rabbit α-VP1 (1:2000, (Rodrigues et al., 1999)), rabbit α-HA (1:1000, Sigma) and rabbit α-myc (1:1000, Sigma). Secondary antibodies were goat α-rabbit IgG, goat α-rat IgG, goat α-mouse IgG conjugated with fluorophores Alexa Fluor^®^ 488, Alexa Fluor^®^ 594 (1:1000, Invitrogen).

### 6.7 Electron microscopy

To prepare samples for electron microscopy non-induced and induced (Day 6) PCF trypanosomes were pelleted by centrifugation (3325rcf/5min), washed once in PBS and submerged with a fixative prepared as follows: 2.5% glutaraldehyde (Agar Scientific, Stansted) in 0.15 M HEPES (Fluka) with an osmolarity of 684 mOsm and adjusted to a pH of 7.41. The cells remained in the fixative at 4°C for at least 24 hours before being further processed as previously described by (Trikin et al., 2016). The images were analyzed with ImageJ.

## 7 Acknowledgements

We would like to thank Roberto Docampo for the VP1 antibody. Christine Clayton for providing the pHD2169 vector. Anneliese Hoffmann for critically reading the manuscript. Electron microscopy sample preparation and imaging were performed with devices supported by the Microscopy Imaging Center (MIC) of the University of Bern.

## 8 Author contributions

BJ and TO designed the experiments.

BJ and SA performed the experiments. BJ performed the bioinformatics analysis, characterized the TbTCTP expression pattern and mechanisms in PCF and BSF trypanosomes, expressed and purified the recombinant TCTP for the antibody production, created all the transgenic cell lines and characterized the phenotype of TCTP downregulation in PCF and BSF trypanosomes. SA acquired STED and TEM images and participated in the analysis of the PCF phenotype upon TCTP RNAi.

BJ and TO wrote the manuscript. SA read and participated in the corrections of the initial draft.

## References

Acunzo, J., Baylot, V., So, A., Rocchi, P., 2014. TCTP as therapeutic target in cancers. Cancer Treat. Rev. 40, 760–769.

Berkowitz, O., Jost, R., Pollmann, S., Masle, J., 2008. Characterization of TCTP, the translationally controlled tumor protein, from Arabidopsis thaliana. Plant Cell 20, 3430–3447.

Bommer, U.-A., Borovjagin, A. V, Greagg, M. a, Jeffrey, I.W., Russell, P., Laing, K.G., Lee, M., Clemens, M.J., 2002. The mRNA of the translationally controlled tumor protein P23/TCTP is a highly structured RNA, which activates the dsRNA-dependent protein kinase PKR. RNA 8, 478–496.

Bommer, U.-A., Thiele, B.-J., 2004. The translationally controlled tumour protein (TCTP). Int. J. Biochem. Cell Biol. 36, 379–385.

Brun, R., Schönenberger, 1979. Cultivation and in vitro cloning or procyclic culture forms of Trypanosoma brucei in a semi-defined medium. Short communication. Acta Trop. 36, 289–92.

Cao, B., Lu, Y., Chen, G., Lei, J., 2010. Functional characterization of the translationally controlled tumor protein (TCTP) gene associated with growth and defense response in cabbage. Plant Cell. Tissue Organ Cult. 103, 217–226.

Chan, T.H.M., Chen, L., Guan, X.Y., 2012a. Role of translationally controlled tumor protein in cancer progression. Biochem. Res. Int. 2012.

Chan, T.H.M., Chen, L., Liu, M., Hu, L., Zheng, B.J., Poon, V.K.M., Huang, P., Yuan, Y.F., Huang, J.D., Yang, J., Tsao, G.S.W., Guan, X.Y., 2012b. Translationally controlled tumor protein induces mitotic defects and chromosome missegregation in hepatocellular carcinoma development. Hepatology 55, 491–505.

Chen, S.H., Wu, P.-S., Chou, C.-H., Yan, Y.-T., Liu, H., Weng, S.-Y., Yang-Yen, H.-F., 2007. A knockout mouse approach reveals that TCTP functions as an essential factor for cell proliferation and survival in a tissue-or cell type-specific manner. Mol. Biol. Cell 18, 2525–32.

Chitpatima, S.T., Makrides, S., Bandyopadhyay, R., Brawerman, G., 1988. Nucleotide sequence of a major messenger RNA for a 21 kilodalton polypeptide that is under translational control in mouse tumor cells. Nucleic Acids Res. 16, 2350.

Clayton, C., 2013. The Regulation of Trypanosome Gene Expression by RNA-Binding Proteins. PLoS Pathog. 9, e1003680.

Diraison, F., Hayward, K., Sanders, K.L., Brozzi, F., Lajus, S., Hancock, J., Francis, J.E., Ainscow, E., Bommer, U. a, Molnar, E., Avent, N.D., Varadi, a, 2011. Translationally controlled tumour protein (TCTP) is a novel glucose-regulated protein that is important for survival of pancreatic beta cells. Diabetologia 54, 368–79.

Docampo, R., Moreno, S.N.J., 2011. Acidocalcisomes. Cell Calcium.

Gachet, Y., Tournier, S., Lee, M., Lazaris-Karatzas, a, Poulton, T., Bommer, U. a, 1999. The growth-related, translationally controlled protein P23 has properties of a tubulin binding protein and associates transiently with microtubules during the cell cycle. J. Cell Sci. 112 ( Pt 8, 1257–1271.

García-Salcedo, J.A., Pérez-Morga, D., Gijón, P., Dilbeck, V., Pays, E., Nolan, D.P., 2004. A differential role for actin during the life cycle of Trypanosoma brucei. EMBO J. 23, 780–9.

Gnanasekar, M., Dakshinamoorthy, G., Ramaswamy, K., 2009. Translationally controlled tumor protein is a novel heat shock protein with chaperone-like activity. Biochem. Biophys. Res. Commun. 386, 333–7.

Gnanasekar, M., Ramaswamy, K., 2007. Translationally controlled tumor protein of Brugia malayi functions as an antioxidant protein. Parasitol. Res. 101, 1533–1540.

Haghighat, N.G., Ruben, L., 1992. Purification of novel calcium binding proteins from Trypanosoma brucei: properties of 22-, 24-and 38-kilodalton proteins. Mol. Biochem. Parasitol. 51, 99–110.

Haile, S., Papadopoulou, B., 2007. Developmental regulation of gene expression in trypanosomatid parasitic protozoa. Curr. Opin. Microbiol. 10, 569–577.

Hammarton, T.C., 2007. Cell cycle regulation in Trypanosoma brucei. Mol. Biochem. Parasitol. 153, 1–8.

Hehl, A., Vassella, E., Braun, R., Roditi, I., 1994. A conserved stem-loop structure in the 3’ untranslated region of procyclin mRNAs regulates expression in Trypanosoma brucei. Proc. Natl. Acad. Sci. U. S. A. 91, 370–4.

Hemphill, A., Lawson, D., Seebeck, T., 1991. The cytoskeletal architecture of Trypanosoma brucei. J. Parasitol. 77, 603–12.

Hinojosa-Moya, J., Xoconostle-Cázares, B., Piedra-Ibarra, E., Méndez-Tenorio, A., Lucas, W.J., Ruiz-Medrano, R., 2008. Phylogenetic and structural analysis of translationally controlled tumor proteins. J. Mol. Evol. 66, 472–483.

Hirumi, H., Hirumi, K., 1989. Continuous cultivation of Trypanosoma brucei blood stream forms in a medium containing a low concentration of serum protein without feeder cell layers. J. Parasitol. 75, 985–9.

Hotz, H.R., Hartmann, C., Huober, K., Hug, M., Clayton, C., 1997. Mechanisms of developmental regulation in Trypanosoma brucei: A polypyrimidine tract in the 3’-untranslated region of a surface protein mRNA affects RNA abundance and translation. Nucleic Acids Res. 25, 3017–3025.

Hsu, Y.-C., Chern, J.J., Cai, Y., Liu, M., Choi, K.-W., 2007. Drosophila TCTP is essential for growth and proliferation through regulation of dRheb GTPase. Nature 445, 785–8.

Jung, J., Kim, M., Kim, M.-J., Kim, J., Moon, J., Lim, J.-S., Kim, M., Lee, K., 2004. Translationally controlled tumor protein interacts with the third cytoplasmic domain of Na,K-ATPase alpha subunit and inhibits the pump activity in HeLa cells. J. Biol. Chem. 279, 49868–75.

Kim, M., Jung, Y., Lee, K., Kim, C., 2000. Identification of the calcium binding sites in translationally controlled tumor protein. Arch. Pharm. Res. 23, 633–6.

Klein, K.G., Olson, C.L., Engman, D.M., 1995. Mitochondrial heat shock protein 70 is distributed throughout the mitochondrion in a dyskinetoplastic mutant of Trypanosoma brucei. Mol. Biochem. Parasitol. 70, 207–9.

Kohl, L., Sherwin, T., Gull, K., 1999. Assembly of the paraflagellar rod and the flagellum attachment zone complex during the Trypanosoma brucei cell cycle. J. Eukaryot. Microbiol. 46, 105–109.

Langdon, J.M., Vonakis, B.M., MacDonald, S.M., 2004. Identification of the interaction between the human recombinant histamine releasing factor/translationally controlled tumor protein and elongation factor-1 delta (also known as eElongation factor-1B beta). Biochim. Biophys. Acta - Mol. Basis Dis. 1688, 232–236.

MacGregor, P., Matthews, K.R., 2012. Identification of the regulatory elements controlling the transmission stage-specific gene expression of PAD1 in Trypanosoma brucei. Nucleic Acids Res. 40, 7705–7717.

Mak, C.H., Su, K.W., Ko, R.C., 2001. Identification of some heat-induced genes of Trichinella spiralis. Parasitology 123, 293–300.

Mani, J., Desy, S., Niemann, M., Chanfon, A., Oeljeklaus, S., Pusnik, M., Schmidt, O., Gerbeth, C., Meisinger, C., Warscheid, B., Schneider, A., 2015. Mitochondrial protein import receptors in Kinetoplastids reveal convergent evolution over large phylogenetic distances. Nat. Commun. 6, 6646.

McKean, P.G., 2003. Coordination of cell cycle and cytokinesis in Trypanosoma brucei. Curr. Opin. Microbiol. 6, 600–607.

Moreno, S.N., Docampo, R., Vercesi, A.E., 1992. Calcium homeostasis in procyclic and bloodstream forms of Trypanosoma brucei. Lack of inositol 1,4,5-trisphosphate-sensitive Ca2+ release. J. Biol. Chem. 267, 6020–6.

Oberholzer, M., Morand, S., Kunz, S., Seebeck, T., 2006. A vector series for rapid PCR-mediated C-terminal in situ tagging of Trypanosoma brucei genes. Mol. Biochem. Parasitol. 145, 117–20.

Portman, N., Gull, K., 2014. Identification of paralogous life-cycle stage specific cytoskeletal proteins in the parasite Trypanosoma brucei. PLoS One 9, e106777.

Pusnik, M., Schmidt, O., Perry, A.J., Oeljeklaus, S., Niemann, M., Warscheid, B., Lithgow, T., Meisinger, C., Schneider, A., 2011. Mitochondrial preprotein translocase of trypanosomatids has a bacterial origin. Curr. Biol. 21, 1738–43.

Rid, R., Önder, K., Trost, A., Bauer, J., Hintner, H., Ritter, M., Jakab, M., Costa, I., Reischl, W., Richter, K., Macdonald, S., Jendrach, M., Bereiter-Hahn, J., Breitenbach, M., 2010. H 2 O 2 ‐dependent translocation of TCTP into the nucleus enables its interaction with VDR in human keratinocytes: TCTP as a further module in calcitriol signalling. J. Steroid Biochem. Mol. Biol. J. Steroid Biochem. Mol. Biol. 118, 29–40.

Rinnerthaler, M., Jarolim, S., Heeren, G., Palle, E., Perju, S., Klinger, H., Bogengruber, E., Madeo, F., Braun, R.J., Breitenbach-Koller, L., Breitenbach, M., Laun, P., 2006. MMI1 (YKL056c, TMA19), the yeast orthologue of the translationally controlled tumor protein (TCTP) has apoptotic functions and interacts with both microtubules and mitochondria. Biochim. Biophys. Acta 1757, 631–8.

Robinson, D.R., Sherwin, T., Ploubidou, A., Byard, E.H., Gull, K., 1995. Microtubule polarity and dynamics in the control of organelle positioning, segregation, and cytokinesis in the trypanosome cell cycle. J. Cell Biol. 128, 1163–72.

Rodrigues, C.O., Scott, D.A., Docampo, R., 1999. Characterization of a vacuolar pyrophosphatase in Trypanosoma brucei and its localization to acidocalcisomes. Mol. Cell. Biol. 19, 7712–23.

Sanchez, J.C., Schaller, D., Ravier, F., Golaz, O., Jaccoud, S., Belet, M., Wilkins, M.R., James, R., Deshusses, J., Hochstrasser, D., 1997. Translationally condrolled tumor protein: A protein identified in several nontumoral cells including erythrocytes. Electrophoresis 18, 150–155.

Sasse, R., Gull, K., 1988. Tubulin post-translational modifications and the construction of microtubular organelles in Trypanosoma brucei. J. Cell Sci. 90 ( Pt 4), 577–89.

Sheffield, P., Garrard, S., Derewenda, Z., 1999. Overcoming expression and purification problems of RhoGDI using a family of &quot;parallel&quot; expression vectors. Protein Expr. Purif. 15, 34–9.

Sherwin, T., Gull, K., 1989. Visualization of detyrosination along single microtubules reveals novel mechanisms of assembly during cytoskeletal duplication in trypanosomes. Cell 57, 211–221.

Stuart, K., Brun, R., Croft, S., Fairlamb, A., Gürtler, R.E., McKerrow, J., Reed, S., Tarleton, R., 2008. Kinetoplastids: related protozoan pathogens, different diseases. J. Clin. Invest. 118, 1301–10.

Thiele, H., Berger, M., Skalweit, a, Thiele, B.J., 2000. Expression of the gene and processed pseudogenes encoding the human and rabbit translationally controlled tumour protein (TCTP). Eur. J. Biochem. 267, 5473–81.

Trikin, R., Doiron, N., Hoffmann, A., Haenni, B., Jakob, M., Schnaufer, A., Schimanski, B., Zuber, B., Ochsenreiter, T., 2016. TAC102 Is a Novel Component of the Mitochondrial Genome Segregation Machinery in Trypanosomes. PLoS Pathog. 12, 1–27.

Tuynder, M., Susini, L., Prieur, S., Besse, S., Fiucci, G., Amson, R., Telerman, A., 2002. Biological models and genes of tumor reversion: cellular reprogramming through tpt1/TCTP and SIAH-1. Proc. Natl. Acad. Sci. U. S. A. 99, 14976–14981.

Vickerman, K., 1965. Polymorphism and Mitochondrial Activity In Sleeping Sickness Trypanosomes. Nature 208, 762–766.

Vickerman, K., 1976. The Diversity of the kinetoplastid flagellatesle. Biol. Kinetoplastida 1–34.

Wheeler, R.J., Scheumann, N., Wickstead, B., Gull, K., Vaughan, S., 2013. Cytokinesis in trypanosoma brucei differs between bloodstream and tsetse trypomastigote forms: Implications for microtubule-based morphogenesis and mutant analysis. Mol. Microbiol. 90, 1339–1355.

Wirtz, E., Leal, S., Ochatt, C., Cross, G.A., 1999. A tightly regulated inducible expression system for conditional gene knock-outs and dominant-negative genetics in Trypanosoma brucei. Mol. Biochem. Parasitol. 99, 89–101.

Wirtz, E., Leal, S., Ochatt, C., Cross, G.A.M., 1999. A tightly regulated inducible expression system for conditional gene knock-outs and dominant-negative genetics in Trypanosoma brucei. Mol. Biochem. Parasitol. 99, 89–101.

Woodward, R., Gull, K., 1990. Timing of nuclear and kinetoplast DNA replication and early morphological events in the cell cycle of Trypanosoma brucei. J. Cell Sci. 95.

